# DeepPHiC: Predicting promoter-centered chromatin interactions using a novel deep learning approach

**DOI:** 10.1101/2022.05.24.493333

**Authors:** Aman Agarwal, Li Chen

## Abstract

**Motivation:** Promoter-centered chromatin interactions, which include promoter-enhancer and promoter-promoter interactions, are important to decipher gene regulation and disease mechanisms. The development of next generation sequencing technologies such as promoter capture Hi-C (pcHi-C) leads to the discovery of promoter-centered chromatin interactions. However, pcHi-C experiments are expensive and thus may be unavailable for tissues or cell types of interest. In addition, these experiments may be underpowered due to insufficient sequencing depth or various artifacts, which results in a limited finding of interactions.

**Results:** To overcome these challenges, we develop a supervised multi-modal deep learning model, which utilizes a comprehensive set of features including genomic sequence, epigenetic signal and anchor distance to predict tissue/cell type-specific genome-wide promoter-enhancer and promoter-promoter interactions. We further extend the deep learning model in a multi-task learning and a transfer learning framework. We demonstrate that the proposed approach outperforms state-of-the-art deep learning methods and is robust to the inclusion of anchor distance as a feature. In addition, we find that the proposed approach can achieve comparable prediction performance using biologically relevant tissues/cell types compared to using all tissues/cell types especially for predicting promoter-enhancer interactions.

**Availability:** https://github.com/lichen-lab/DeepPHiC

## 1 Introduction

Three-dimensional chromatin interactions via chromatin looping play a key role in gene regulation, which allows distal regulatory regions (e.g., enhancer) to be physical proximity to their target gene promoters to regulate transcription. However, the distance of the two regions on chromosome can be tens of kilobases (kb) away from each other. Various experimental approaches have been developed for detecting chromatin interactions, which can be captured at the level of a single locus by low-throughput 3C, 4C, a set of loci by medium-throughput 5C, ChIA-PET and genome-wide loci by high-throughput *in situ* Hi-C [1]. However, 3C and 4C are not practical for a genome-wide interrogation, and ChIA-PET can only identify interactions involving highly transcriptionally active genes but not for long-range interactions involving weakly expressed and transcriptionally inactive genes. The recent developed high-throughput *in situ* Hi-C can simultaneously interrogate genome-wide chromatin interactions.

Promoter-enhancer interactions, where enhancer initializes or activates the level of transcription of the targeted gene, is well studied. Promoter-promoter interactions, which have been reported in cultured mammalian cells and a few primary tissues, also play an important role in transcriptional coordination in the way that proximal and distal genes are active and transcribed cooperatively through promoter-promoter interactions [2, 3, 4]. To interrogate both promoter-enhancer interactions and promoter-promoter interactions, promoter-centered capture Hi-C (pcHi-C) has emerged as a cost-effective method to map chromatin interactions for promoter regions at a high resolution by targeting promoters directly. However, it is costly to sequence deeply for a high spatial resolution to interrogate promoter-centered chromatin interactions. Thus, pcHi-C experiments are expensive and may not exist for tissues or cell types of interest. In addition, pcHi-C experiments may be underpowered due to insufficient sequencing depth or various artifacts, which results in a limited finding of interactions. Therefore, it is demanded to develop a computational model to accurately predict tissue/cell type-specific promoter-centered chromatin interactions even in unknown tissues.

The advent and advance of deep learning provides an unprecedented opportunity to derive training features and nonlinear dependencies from the genomic sequence. Towards this direction, deep learning has been widely utilizes to predict various epigenetic signals such as transcription factor binding site [5], chromatin accessibility [6] and DNA methylation [7]. Recently, supervised deep learning methods have been widely utilized to perform *in silico* prediction of genome-wide chromatin interactions. The general principle of these methods is to treat genomic sequences of two anchors as the model input and predict the probability of the anchor interaction, while these methods differ from the neural network architecture, the training set and the input features. For example, SPEID is a simple convolutional neural network (CNN), which uses DNA sequence in two anchors to predict promoter-enhancer interactions [8]. SPEID consists of two subnetworks, one for each anchor, to extract sequence features using convolutional layers and merge the extracted features into a long short-term memory network (LSTM) layer before making the prediction. SPEID is trained using labeled chromatin interactions derived from *in situ* Hi-C data in six cell lines (e.g, GM12878, HeLa-S3, IMR90, K562) [9]. DeepMILO adopts a combination of a CNN and a bidirectional LSTM to predict the insulator loops by using labeled chromatin interactions derived from ChIA-PET data in four cell lines including GM12878, K562, MCF7, and Hela3 [10]. ChINN is a multi-modal deep learning model, which uses both the genomic sequence of two anchors and the distance between the two anchors to predict the chromatin interactions [11]. The neural network architecture of ChINN is modified based on DeepSEA [5]. Particularly, ChINN demonstrates that inclusion of anchor distance as a feature improves the prediction performance. In addition, ChINN is trained and evaluated on ChIA-PET datasets, *in situ* Hi-C datasets from chronic lymphocytic leukemia patient samples, and *in situ* Hi-C datasets from Rao. et al. [9]. DeepTACT is also a multi-modal deep learning model. Different from ChINN, DeepTACT [12] integrates genomic sequence and chromatin accessibility data in the anchors to predict both promoter–enhancer interactions and promoter–promoter interactions by using promoter capture Hi-C (pcHi-C) datasets from B cell and CD cell lines [3].

Despite the success of the previous work, there are still several challenges. First, to our best knowledge, there is no deep learning approach yet to include genomic sequence and epigenetic signal in the anchors as well as anchor distance into an integrative framework. Second, deep learning approaches, which target to predict tissue/cell type-specific promoter-promoter and promoter-enhancer interactions in a broad context, is underdeveloped. Third, the sample size of labelled chromatin interactions can be moderate, which makes the the training set less powerful for achieving a genome-wide prediction. In this work, we develop a multimodal deep learning framework named DeepPHiC, which is based on the densely connected convolutional neural network, to predict both promoter-promoter and promoter-enhancer interactions. The contribution of our work lies on several aspects: (i) DeepPHiC is trained and evaluated on a comprehensive set of pcHi-C data including 27 tissues and cell types; (ii) DeepPHiC utilizes a comprehensive set of informative features, ranging from genomic sequence, epigenetic signal in the anchors and anchor distance; (iii) An extension of DeepPHiC to a multi-task learning and transfer learning framework further improves the prediction performance; (iv) DeepPHiC is released as an open-source python toolkit, which can benefit the genomic research community. As a result, we demonstrate that inclusion of more informative features leads to improved prediction performance; the multi-task learning and transfer learning framework further improve the prediction performance. Importantly, DeepPHiC outperforms state-of-the-art deep learning approaches for the similar purpose and is robust to the inclusion of anchor distance as a feature. Finally, we find that using biologically relevant tissues/cell types for multi-task learning and transfer learning of DeepPHiC can achieve comparable performance to using all tissues/cell types, especially for predicting promoter-enhancer interactions.

## 2 Methods

### 2.1 Tissue/cell type-specific promoter-centered chromatin interactions

We collect and curate a deposition of promoter-centered chromatin interactions in 3D-genome Interaction Viewer and database (3DIV) [13], which has profiled a comprehensive set of promoter-centered capture Hi-C datasets (pcHi-C) across 27 human cell/tissue types in 4 categories including embryonic stem cell, 4 early embryonic lineages, 2 primary cell lines and 20 primary tissue types, which also has corresponding epigenome datasets in Roadmap Epigenomics Project [14]. For each tissue/cell type, pcHi-C provides both promoter-promoter interactions (PP) and promoter-enhancer interactions (PE).

To create the labelled training set, we filter chromatin interactions with distance more than 10^6^ and consider only intra-chromosomal chromatin interactions. Then, we format each anchor into 2kb region with 1kb upstream and downstream of the midpoint of the anchor. Among all interactions, we label significant interactions (FDR*<*0.1) as the positive set and only keep tissues/cell types with more than 100 significant interactions. As a result, the number of positive samples ranges from 972 to 56057 with a median of 5071 among 26 tissues/cell types for PE and from 187 to 10338 with a median of 917 among 25 tissues/cell types for PP. To create the negative set, we select interactions (FDR*>*0.5) with matched GC contents as the positive set. Without loss of generality, the number of negative interactions is set as the same as the positive interactions. More details of the distribution of positive interactions in tissues/cell types can be found in Table S1 and Table S2.

### 2.2 Tissue/cell type-specific epigenetic datasets

To create the epigenetic signals in the anchors, we collect the matched tissue/cell type-specific epigenetic datasets from Roadmap Epigenomics Project [14]. Especially, we collect open chromatin data and active histone marks associated with promoter activation such as H3K4me3, and active histone marks associated with enhancer activation such as H3K4me1 and H3K27ac because these histone marks have been suggested important to the formation of chromatin interactions and used for predicting chromatin interactions [15]. More details of matched epigenetic datasets for pcHi-C datasets can be found in the Table S3.

### 2.3 Multi-modal features of DeepPHiC

DeepPHiC consists of three types of input features, which include genomic sequence and epigenetic signal in the anchors as well as anchor distance. Specifically, we obtain the 2000bp genomic sequence for each anchor by extending the middle of each anchor upstream and downstream 1000bp. We further perform one-hot encoding for the genomic sequence by following the rule as ‘A’: [1,0,0,0], ‘C’: [0,1,0,0], ‘G’: [0,0,1,0] and ‘T’: [0,0,0,1]. As a result, the genomic sequence is converted into a 2000 × 4 matrix. Moreover, anchor distance is defined as the distance between the middles of two anchors. For the epigenetic feature, we obtain the matched tissue/cell type-specific epigenetic datasets from Roadmap Epigenomics Project [14], and calculate the normalized read counts in a window size of 1000 with a sliding size 50, yielding 21 read features for each anchor.

### 2.4 Network architecture of DeepPHiC

The network architecture of DeepPHiC is developed based on the DenseNet [16] because densely connected neural network has been demonstrated to promote feature reuse and therefore reduce the number of parameters. Similar to DenseNet, DeepPHiC adopts a ResNet-style structure with skip connections [17], wherein previous layers are connected to all subsequent layers in the network via feature concatenation. As a result, during back propagation, each layer has a direct access to the output gradients, resulting in faster network convergence and regularization.

DeepPHiC consists two modules: feature extractor module and classifier module. The feature extractor module consists of two dense blocks and one convolutional layer outside of the dense blocks. A convolution layer in DeepPHiC is defined as a sequence of four operations: convolution, batch normalization [18], PReLU non-linearity [19], and dropout [20]. A dense block in DeepPHiC is made up of four such convolution layers with different number of filters and dilation rates. The sequence feature and read feature are passed separately through their own dense block to produce high-level features, which are concatenated and connected to subsequent convolution layer outside of the dense block for generating the fused features. The fused features are then concatenated to the distance feature to produce the extracted features. The extracted features are fed to the classifier module, which only contains a fully-connected layer for the final prediction. Specifically, since the dimensions of sequence feature and read feature are different, we use different kernel sizes and strides accordingly. For sequence feature, the kernel size is 20 × 4 with a stride of 10 × 4, whereas, for read feature, the kernel size is 3 × 1 with stride of 1 × 1. For both features, we use 64, 32, 16 and 16 filters and dilation rates of 1, 2, 3, and 4 respectively in the four convolutional layers in the dense block. The stride for last three convolution layers in the dense block were 1 × 1 so that the features are made the similar dimension for concatenation. The convolution layer outside of the dense blocks has the kernel size 4 × 4 and stride 4 × 4, which is used for feature fusion. This layer is followed by global average pooling layer and is concatenated with the distance feature, which are then passed to a fully-connected layer with 256 neurons and activated by ReLU. The output layer with the Sigmoid activation function predict the probability of two anchors are interacted.

### 2.5 Multi-task learning paradigm of DeepPHiC

In the multi-task learning framework, we define the prediction for chromatin interaction in each tissue as a single task, and the prediction of chromatin interaction in multiple tissues simultaneously forms a multitask learning paradigm. A common practice of multi-task learning model, such as DeepSEA, is a fully shared model. DeepSEA is essentially a deep convolutional neural network with multi-label outputs [5]. It takes genomic sequence as the model input to predict multiple epigenetic signals from thousands of tissues simultaneously. All the layers in the network are shared by the prediction tasks for all tissues. However, the fully shared model may not be ideal for several reasons. First, chromatin interactions can be overlapped among tissues, which indicate common features shared among tissues. Moreover, chromatin interactions can also be unique to certain tissues, indicating tissue-specific private features. Second, since features learned by the fully shared model contains both common features and private features, it is difficult to distinguish common features from private features. Therefore, common features are prone to be contaminated by private features, and vice versa. In addition, private features from different tissues will interfere with each other, that is, private features of dominated tissues with a large sample size, will adversely affect the prediction of other tissues with a small sample size. These unfavorable bias will compromise the goal of multi-task learning. Therefore, it is important to disentangle common features shared by all tissues from tissue-specific private features by developing a more advanced multi-task neural network architecture.

Due to the above considerations, we adapt the multi-task learning paradigm [21, 22] to develop the multi-task DeepPHiC, which contains a shared feature extractor (Figure 1A), to learn common features for all tissues, and a private feature extractor to learn tissue-specific private features. The integrated features from both the common features and private features will connect to the tissue-specific classifier for predicting tissue-specific chromatin interactions. The disentanglement of private features from common features has the advantage to alleviate the interference among private features from multiple tissues, and between common features and private features. Moreover, multi-task DeepPHiC will likely improve the prediction performance for chromatin interactions in tissues with a moderate sample size, by leveraging labelled chromatin interactions from other tissues to correctly learn the common features. Specifically, to train the multi-task DeepPHiC, we will pretrain a fully-shared feature extractor by pooling labelled chromatin interactions from all tissues except the tissue of interest to learn the common features. Next, labelled chromatin interactions from the tissue of interest will be used to train the tissue-specific feature extractor. Then, the fully-shared feature extractor will be transferred into the shared-private model (Figure 1B).

**Figure 1:**
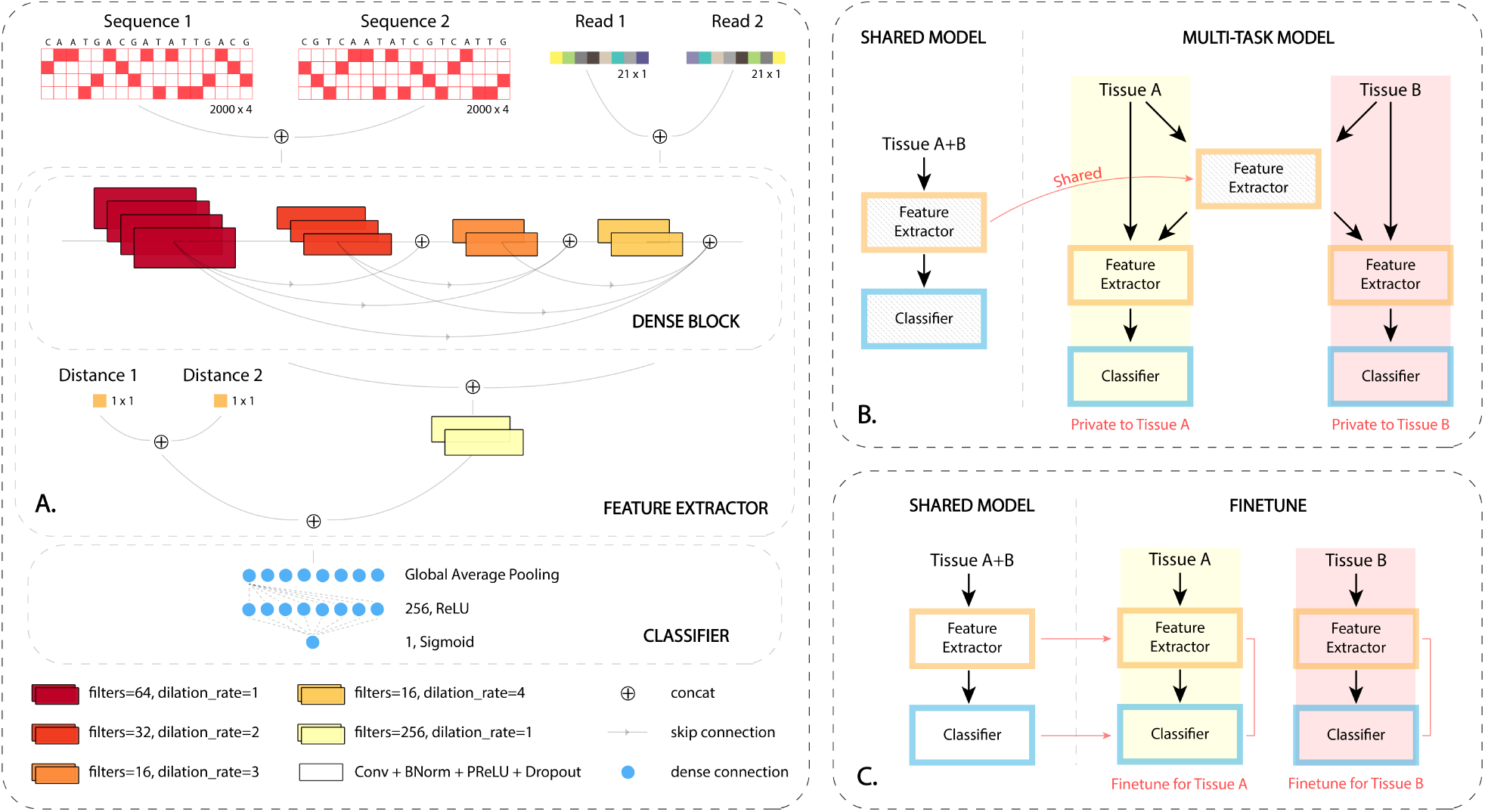
Overview of DeepPHiC. **A**. Model architecture of DeepPHiC. **B**. Multi-task paradigm of Deep-PHiC (DeepPHiC-ML). In practice, when tissue A is of interest, we aggregate all chromatin interactions from other tissues except tissue A to pretrain the shared feature extractor. Next, we will use the chromatin interactions from tissue A to train the tissue-specific feature extractor. The final prediction is obtained by integrating common features obtained from the shared feature extractor and tissue-specific private features obtained from the tissue-specific feature extractor to the tissue-specific classifier. Similarly, if tissue B is of interest, all chromatin interactions from other tissues except tissue B are used to pretrain the shared feature extractor. Thus, for each tissue, there is an independent multi-task model. **C**. Transfer learning paradigm of DeepPHiC. All tissues except the tissue of interest is used to pretrain the shared feature extractor. For each tissue, fine-tuning of DeepPHiC is performed for both shared feature extractor and tissue-specific classifier. Thus, for each tissue, there is an independent transfer learning model. For both DeepPHiC-ML and DeepPHiC-TL, 85% labelled data is used for training and 15% for validation in pretraining the shared feature extractor, while 70% labelled data is used for training, 15% for validation in fine-tuning the whole network and 15% for independent testing the whole network.

### 2.6 Transfer learning paradigm of DeepPHiC

Transfer learning approach has been widely used for improving the prediction for targeted task by leveraging information from a source task [23]. A common assumption for transfer learning is that the feature space in source task is similar to the target task and therefore both tasks share common features. The common features can be learnt by the source task and transferred to the target task. The target task can be used to fine-tune the task-specific features. Therefore, transfer learning can leverage both labelled data from source task, which usually has a large sample size, and labelled data from the target task with a small sample size to improve the prediction performance for the target task. Particularly, deep transfer learning, which utilizes deep neural network in the transfer learning, has been widely adopted in genomic study [24]. Practically, the first layers are pretrained using labelled data from the source task, frozen and transferred to the target task, where the last layers or all layers are fine-tuned using the labelled data from the target task. Here, we define the prediction for chromatin interactions in any tissue as the source task, and the prediction of chromatin interactions in the tissue of interest as the targeted task. In practice, we aggregate chromatin interactions from all tissues to pretrain the common feature extractor, freeze it and transfer it to each single task (1C). In each single task, the whole network, which includes shared feature extractor and task-specific classifier, is fine-tuned by the tissue-specific features.

### 2.7 Model implementation, training and testing

All three paradigms of DeepPHiC are implemented using TensorFlow [25] on an NVIDIA V100 GPU system. Specifically, the network is optimized for a binary cross-entropy loss function using the mini-batch gradient descent and the Adam optimizer [26]. The initial learning rate is set to 1e-4 with the exponential decay. Each models is trained for a maximum of 200 epochs, and the training is stopped when the model’s loss stopped decreasing for at least five consecutive epochs.

For each tissue, three paradigms of DeepPHiC are trained, which are original DeepPHiC (DeepPHiC-Base), multi-task paradigms of DeepPHiC (DeepPHiC-ML) and transfer learning paradigms of DeepPHiC (DeepPHiC-TL). DeepPHiC-Base is trained for each tissue independently with 70% labelled data for training, 15% for validation, 15% for testing, and no data from other tissues will be used. For DeepPHiC-ML, chromatin interactions from other tissues except tissue of interest are used to pretrain the shared feature extractor. Chromatin interactions from tissue of interest are used to train the tissue-specific feature extractor for obtaining the private features, together with the common features from the shared feature extractor, to train the tissue-specific classifier. For DeepPHiC-TL, all tissues except the tissue of interest is used to pretrain the shared feature extractor, and fine-tuning of DeepPHiC is performed for the whole network. For both DeepPHiC-ML and DeepPHiC-TL, 85% labelled data is used for training and 15% for validation in pretraining the shared feature extractor, 70% labelled data is used for training and 15% for validation in fine-tuning the whole network, and 15% for independent testing the whole network.

### 2.8 Competing methods and evaluation metrics

To demonstrate the superior performance of DeepPHiC, we compare DeepPHiC to four state-of-the-art deep learning methods with the similar purpose to predict chromatin interactions. Specifically, SPEID is convolutional neural network, which only uses genomic sequence in the anchors as the input features. Similarly, DeepMILO also utilizes genomic sequence in the anchors but extends the model by integrating a deep convolutional neural network and a bidirectional long short-term memory network, which will capture both local and long distance sequence dependency. Particularly, ChINN and DeepTACT are two multi-modal deep learning models. ChINN use both anchor distance and genomic sequence in the anchors as the input features. DeepTACT integrates both genomic sequence in the anchors and chromatin accessibility in the anchors as the input features to predict both promoter-enhancer interactions and promoter–promoter interactions. We use both Area Under the Curve (AUC) and Area Under the Precision Recall Curve (AUPRC) to evaluate the prediction performance for all methods. To alleviate the bias of random sampling, for each model, we randomly split the whole labelled data 10 times into training, testing and validation sets, and report AUC an AUPRC for 10 experiments.

## 3 Results

### 3.1 A combination of genomic sequence, epigenetic signal in the anchors and anchor distance improves the prediction for promoter-centered chromatin interactions

Most existing deep learning approaches with the similar purpose adopt the genomic sequence in the anchors as the model input such as SPEID and DeepMILO. Other approaches expand the feature sets by including anchor distance (e.g., ChINN) and epigenetic signal in the anchors (e.g., DeepTACT). Nevertheless, none of these approach considers all three types of features simultaneously. Without loss of generality, we use labelled chromatin interactions from both PE and PP in Lymphoblastoid cells (GM) to demonstrate that integrating all three types of features together will achieve the best prediction performance. Since DeepPHiC is a multi-modal model, which takes sequence (S), read (R) and distance (D) as the model input, we adapt DeepPHiC to make it accommodate to arbitrary combinations of feature sets in the feature extractor accordingly (Figure 1A).

As a result, we find that using all three types of features (S+R+D) achieves the highest AUC and AUPRC in predicting both PE and PP, followed by sequence and distance (S+D), and distance (D) (Figure 2A, B). Interestingly, using distance alone (D) already achieves promising prediction performance, which indicates that promoter-centered interactions are closer in genome than random interactions. Integrating sequence and distance (S+D) does not help much improve the prediction performance. However, combining sequence and read (S+R) outperforms using sequence alone (S). These observations indicate that it is important to consider all three types of features as model input in a multi-modal framework in DeepPHiC. We will use the multi-modal DeepPHiC (S+R+D) as default implementation when the tissue-matched epigenetic data is available. Otherwise, we will use multi-modal DeepPHiC (S+D) since genomic sequence and anchor distance are readily available in the Hi-C summary data.

**Figure 2:**
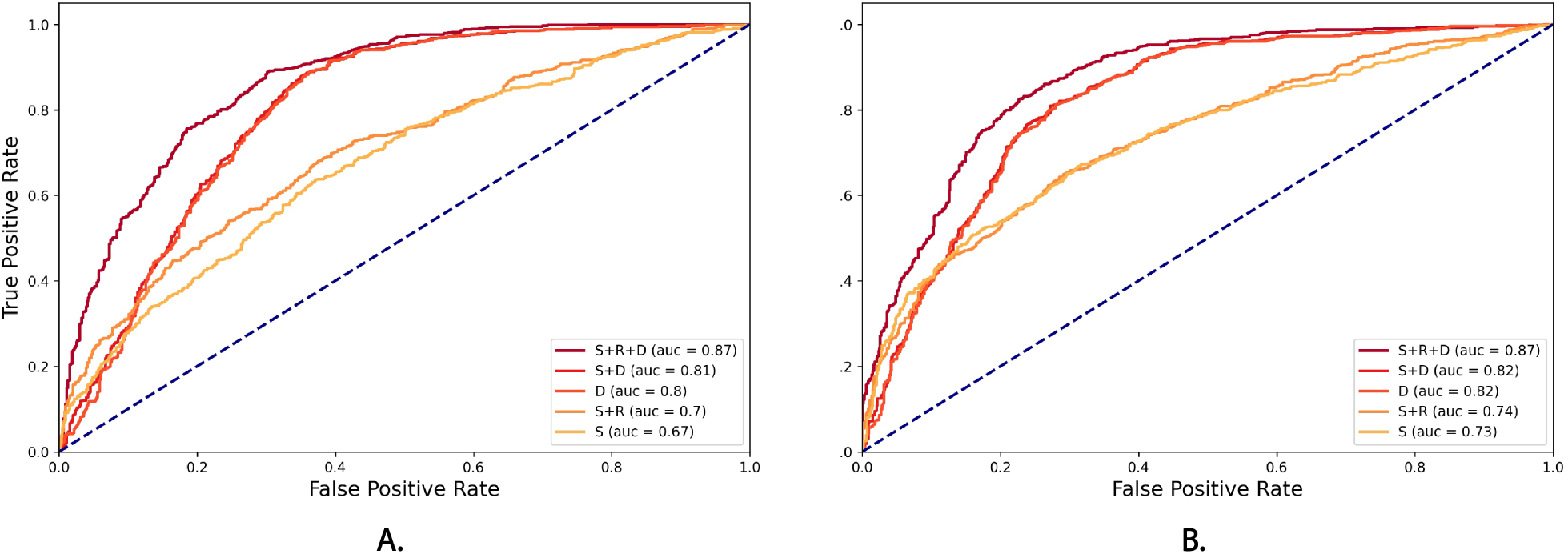
Comparison of performance of different combination of feature sets for Lymphoblastoid cells (GM) (S: sequence, R: read, D: distance) in DeepPHiC. **A**. Promoter-enhancer interaction **B**. Promoter-promoter interaction

### 3.2 Comparison between DeepPHiC-Base, DeepPHiC-ML and DeepPHiC-TL

For each tissue, we train DeepPHiC-Base, which is trained only using labelled chromatin interactions from this tissue. We also train DeepPHiC-ML and DeepPHiC-TL, both of which use labelled data from both tissue of interest and other tissues. We evaluate all three paradigms of DeepPHiC to predict PE in 26 tissues and PP in 25 tissues, and report AUC and AUPRC in 10 experiments respectively.

We use “mAUC” to represent median of AUCs in 10 experiments. For each tissue, we calculate the rank of mAUC for three paradigms of DeepPHiC. For PE prediction, DeepPHiC-TL has the best performance in 22 out of 26 tissues, followed by DeepPHiC-ML, which achieves the second best performance in 17 out of 26 tissues. However, DeepPHiC-Base has the lowest AUC in 16 out of 26 tissues. Similarly, for PP prediction, DeepPHiC-TL has the highest AUC in 23 out of 25 tissues, followed by DeepPHiC-ML, which obtains the second best performance in 21 out of 25 tissues. In contrast, DeepPHiC-Base has the lowest AUC in 21 out of 25 tissues (Figure 3). The above observations indicate that both DeepPHiC-TL and DeepPHiC-ML outperform DeepPHiC-Base by benefiting leveraging labeled chromatin interactions from other tissues. We also measure the prediction performance of all methods by AUPRC and find a similar trend (Figure S1). Since DeepPHiC-TL has the overall best performance, we will use it as the default implementation in the following sections to compare with state-of-the-art deep learning models.

**Figure 3:**
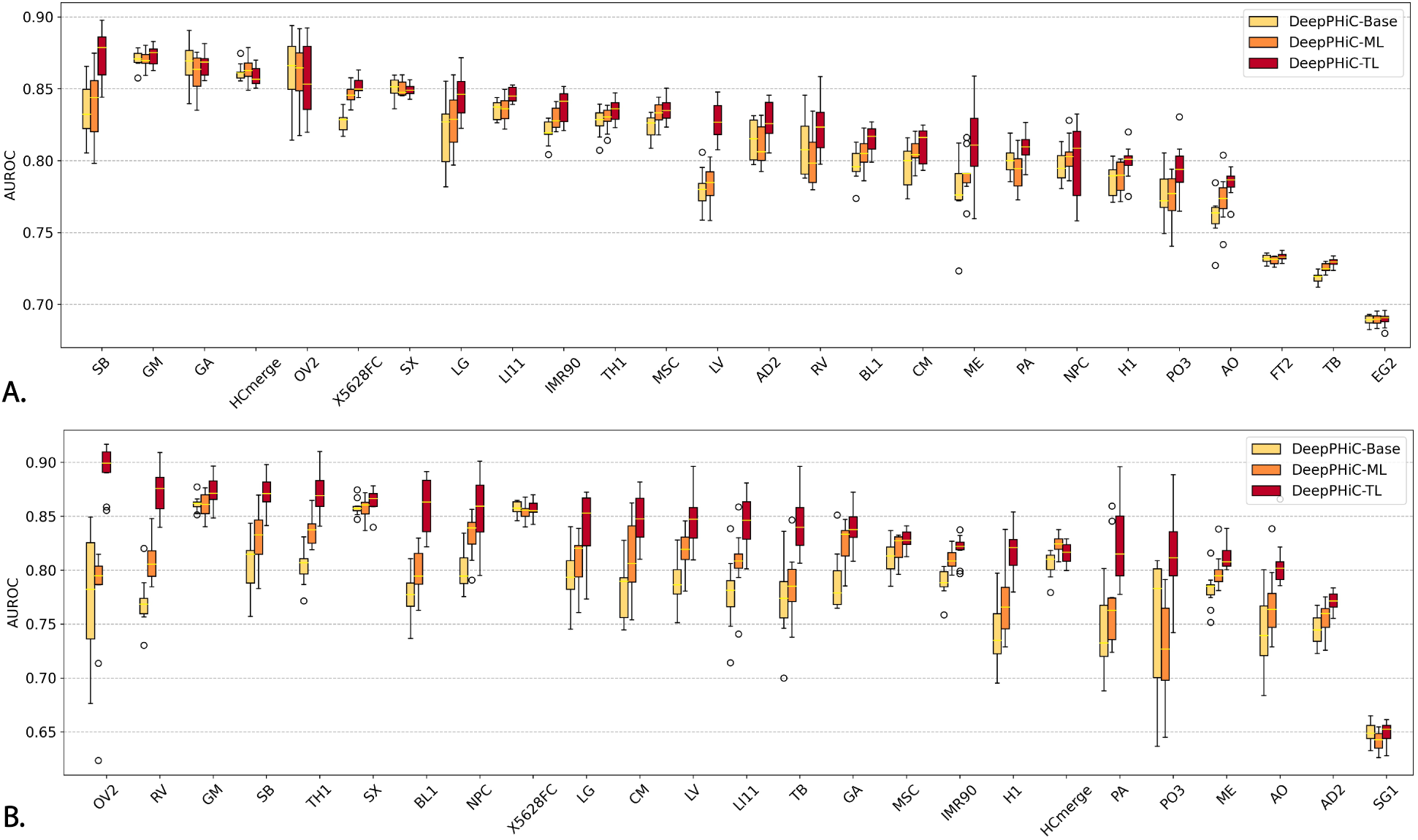
Comparison of three paradigms of DeepPHiC. **A**. promoter-enhancer interactions (PE) in 26 tissues/cell types and **B**. promoter-promoter interactions (PP) in 25 tissues/cell types. AUC are reported on 10 different experiments.

Since both DeepPHiC-ML and DeepPHiC-TL improve the learning for common feature space by lever-aging labelled chromatin interactions from other tissues, the advantage of DeepPHiC-ML and DeepPHiC-TL to DeepPHiC-Base is more evident for tissue with small amount of labeled data. It is because learning common feature space may be underfitting by small amount of labelled data. For PE prediction, the improvement of AUC is more evident for tissues with a small amount of labelled data such as SB (972), LG (1188), LV (1565) and ME (1045), compared to tissues with a large amount of labelled data such as FT2 (56057), TB (83604) and EG2 (20049), where the three paradigms of DeepPHiC show a comparable performance. Similarly, for PP prediction, among tissues with a small amount of labelled data such as OV2 (187), RV (723), SB (704), TH1 (1066), BL1 (555), NPC (546), LG (661), CM (962), LV (968), LI111 (706), TB (444), GA (900), H1 (917), PA (396), AO (788), the improvement of DeepPHiC-ML and DeepPHiC-TL is more apparent compared to tissues with a large sample size such as GM (4446), SX (5866), X5628FC (6224), MSC (4168), IMR90 (4156), HCmerge (2519), ME (3142), AD2 (5421) and SG1 (10338). Overall, these observations indicate that DeepPHiC-ML and DeepPHiC-TL indeed improve the prediction performance for tissues with a small amount of labelled data. For tissues with a large amount of labelled data, DeepPHiC-ML and DeepPHiC-TL are on par with DeepPHiC-Base. Moreover, since PP are much fewer than PO, it is more favorable to apply DeepPHiC-ML and DeepPHiC-TL for predicting PP.

### 3.3 Comparing DeepPHiC and state-of-the-art deep learning models to predict promoter-enhancer interaction and promoter-promoter interaction

Multiple deep learning models, such as DeepTACT, DeepMILO, SPEID and ChINN, have been developed to predict genome-wide chromatin interactions, which vary in the training set, the types of input features and the network architecture. To make a fair comparison between DeepPHiC and these methods, we use the same training, validation and testing sets for all methods, which are derived from labelled PE in 26 tissues/cell types and labelled PP in 25 tissues/cell types. For competing methods, 70% labelled data is used for training, 15% for validation and 15% for testing for each tissue. For DeepPHiC, we use DeepPHiC-TL as it has been demonstrated to outperform DeepPHiC-Base and DeepPHiC-ML.

For each tissue, we use “mAUC” to represent median of AUCs in 10 experiments and calculate the rank of mAUC for each method. For each method, we summarize the ranks across all tissues. Consequently, DeepPHiC achieves the best performance in all 26 tissues for PE prediction, followed by ChINN, which obtains the second best performance in 25 out of 26 tissues. Nevertheless, SPEID, DeepMILO and DeepTACT have less desirable but comparable performance. Among all methods, we further compare the median of mAUCs across all tissues by using two-sided Wilcoxon rank sum test (Figure 4).

**Figure 4:**
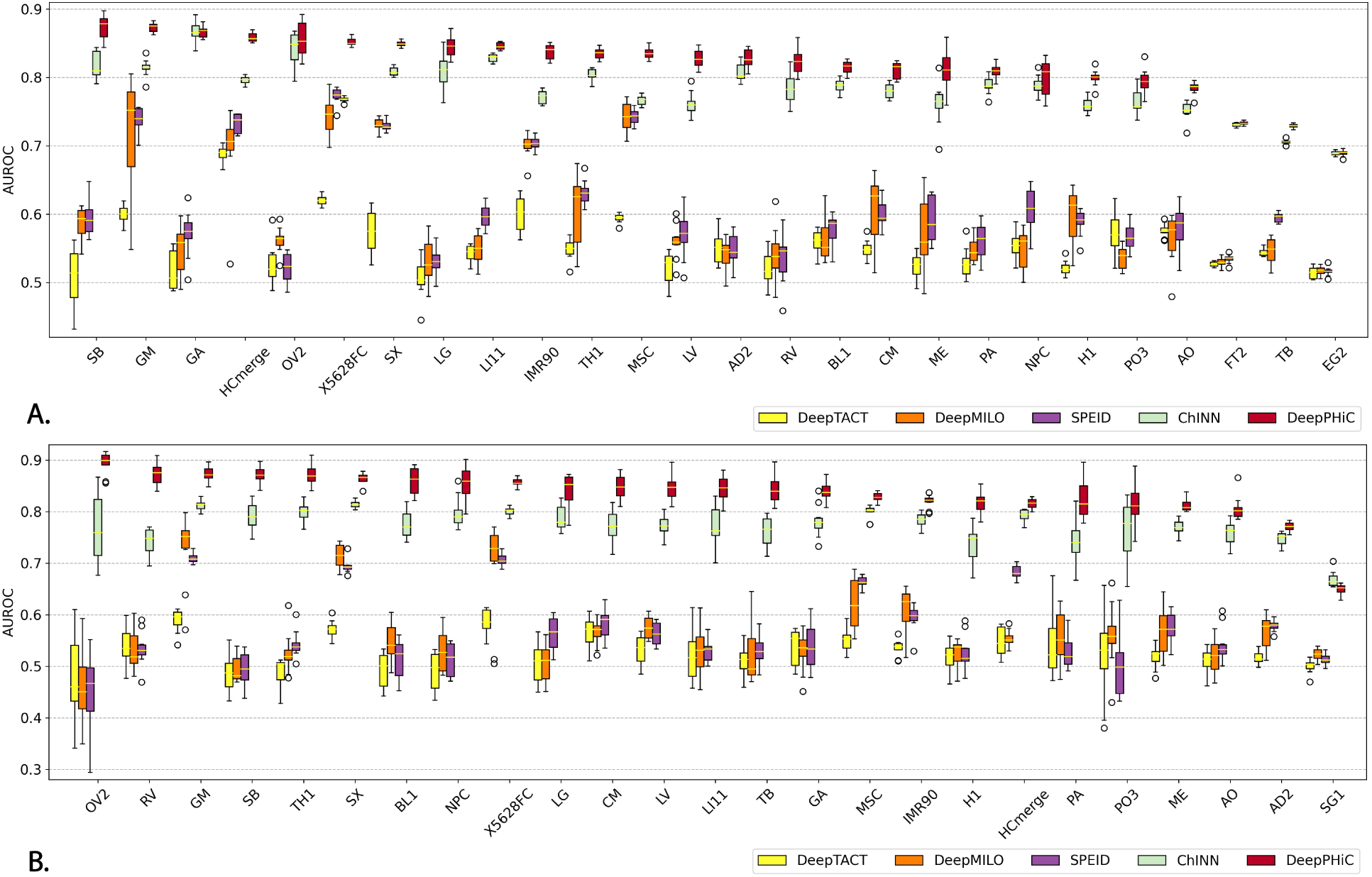
Comparison of DeepPHiC and state-of-the-art deep learning models on predicting **A**. Promoter-enhancer interactions (PE) and **B**. Promoter-promoter interactions (PP). Each boxplot represents the AUC in 10 experiments for each tissue.

As expected, DeepPHiC obtains the highest median of mAUCs for PE prediction, followed by ChINN (DeepPHiC: 0.826; ChINN: 0.785; SPEID: 0.589; DeepMILO: 0.560; DeepTACT: 0.547). The improvement of DeepPHiC over other methods is also significant (pvalue=4.115×10^−4^ for DeepPHiC vs ChINN; pvalue=2.049×10^−12^ for DeepPHiC vs SPEID; pvalue=1.504×10^−12^ for DeepPHiC vs DeepMILO; pvalue=4.033×10^−15^ for DeepPHiC vs DeepTACT). Similarly, DeepPHiC tops other methods in 24 out of 25 tissues for PP prediction, ChINN has the second best performance by ranking second in 24 out of 25 tissues and other approaches have similar performance. Again, DeepPHiC has the highest median of mAUCs, followed by ChINN (DeepPHiC: 0.835; ChINN: 0.744; SPEID: 0.553; DeepMILO: 0.555; DeepTACT: 0.526). The improvement of DeepPHiC over other methods is also significant (pvalue=2.294×10^−9^ for DeepPHiC vs ChINN; pvalue=3.006×10^−13^ for DeepPHiC vs SPEID; pvalue=1.108×10^−13^ for DeepPHiC vs DeepMILO; pvalue=1.582×10^−14^ for DeepPHiC vs DeepTACT). Moreover, we report the prediction performance in terms of AUPRC and find all methods follow a similar trend (Figure S2).

The advantage of DeepPHiC can be explained by several reasons. First, compared to other approaches, it uses the most comprehensive feature types, which include anchor distance, genomic sequence and epigenetic signal in the anchors. Accordingly, DeepPHiC (S+R+D) tops other methods and ChINN (S+D) has the second best performance. This observation is also consistent with the previous finding that S+R+D achieves the best performance, followed by S+D (Figure 2). Moreover, anchor distance is an informative feature as the performance of methods without using anchor distance is significantly declined. Second, DeepPHiC benefits from the transfer learning, which improves the learning for common feature space by leveraging labelled data from other tissues. Since labelled PP are much less than PE (Table S1), transfer learning makes DeepPHiC hold more advantage to ChINN in predicting PP than PE, which is evidenced by a median of 6.6% increased in predicting PP across 26 tissues compared to a median of 3.5% increased in predicting PE across 25 tissues. Specifically, we find that tissues with a moderate size of labelled data can benefit more from transfer learning. For example, compared to ChINN in predicting PP, DeepPHiC improves 13.9% AUC in OV2 (187), 12.7% in RV (723), 9.3% in BL1 (555), 8.4% in LI11 (706), 8.1% in SB (704) and another 12 tissues with more than 5% increase in AUC. In contrast, there is a moderate improvement for tissues with a large size of labelled data such as 2.2% improved AUC for HCmerge (2519) and 1.9% improved AUC for AD2 (5421).

### 3.4 Robustness of DeepPHiC to anchor distance feature

So far, the training set is created based on the confidence of chromatin interactions with matched GC content between the positive set (FDR*<*0.1) and the negative set (FDR*<*0.5). Another way to create the negative set is to match the anchor distance between chromatin interactions in the positive set and the negative set (e.g. DeepTACT). In this case, distance is no longer used as a feature for model input. To demonstrate DeepPHiC is still robust without including anchor distance as a feature, we remove the anchor distance from both DeepPHiC and ChINN and use same aforementioned experimental design to compare all methods in terms of AUC and AUPRC.

As a result, the performance of both ChINN and DeepPHiC declines. For PE prediction (Figure 5A), the median of mAUC of DeepPHiC across all tissues decreases from 0.826 to 0.664 by 16.2%, and the median of mAUC of ChINN across all tissues decreases from 0.785 to 0.541 by 24.4%. Despite the declined performance, DeepPHiC still significantly outperforms other competing approaches (DeepTACT: 0.547; DeepMILO: 0.560; SPEID: 0.589) by obtaining a pvalue less than 0.05 from a two-sided Wilcoxon rank sum test. However, the performance of ChINN declines more compared to DeepPHiC, and ChINN is on par with other approaches. Similarly, for PP prediction (Figure 5B), the median of mAUC of DeepPHiC across all tissues decreases from 0.846 to 0.727 by 11.9%, and the median of mAUC of ChINN across all tissues decreases from 0.773 to 0.551 by 22.2%. Again, DeepPHiC is more robust to ChINN when anchor distance is not considered as the input feature. In addition, DeepPHiC holds a significant advantage over other approaches (DeepTACT: 0.522; DeepMILO: 0.541; SPEID: 0.534) by obtaining a pvalue less than 0.05 from a two-sided Wilcoxon rank sum test. We also report the prediction performance in terms of AUPRC and find all approaches follow a similar trend (Figure S3). Overall, the above findings indicate that DeepPHiC still has the best performance without considering the anchor distance as an input feature.

**Figure 5:**
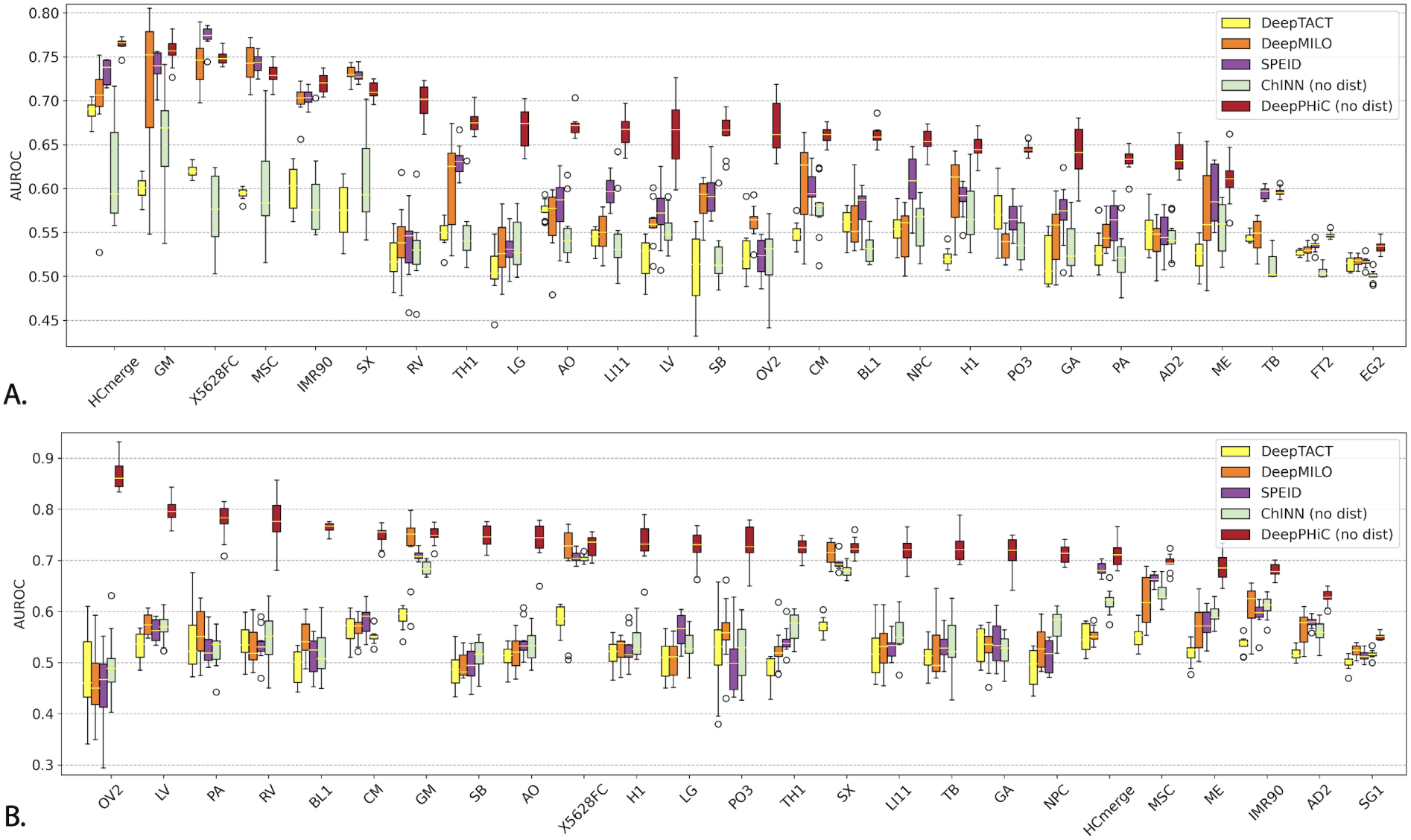
Compare DeepPHiC without distance (DeepPHiC (no dist))and ChINN without distance (ChINN (no dist)) to other deep learning models for predicting **A**. Promoter-enhancer interactions (PE) and **B**. Promoter-promoter interactions (PP). Each boxplot represents the AUC in 10 experiments for each tissue.

### 3.5 Pre-training DeepPHiC with biologically relevant tissues/cell types

Among all tissues/cell types, some tissues/cell types are more relevant to each other (e.g, belonging to the same organ). Hence, we evaluate the prediction performance of DeepPHiC-TL and DeepPHiC-ML by utilizing biologically relevant tissues/cell types. Particularly, we consider three groups of tissues/cell types: (i) Cell types relevant to H1 (H1 group), which include H1 Embryonic Stem Cell (H1), H1-derived Mesendoderm Cell (ME), H1-derived Mesenchymal Stem Cell (MSE) and H1-derived Neuronal Progenitor Cell (NPC); (ii) Brain regions (Brain group), which include Hippocampus (HCmerge) and Dorsorateral prefrontal cortex (X5628FC); (iii) Cell types relevant to heart (Heart group), which include Cardiomyocytes (CM), Aorta (AO), Left Ventricle (LV) and Right Ventricle (RV).

For each group of tissues/cell types, we perform the same aforementioned procedure to train DeepPHiC-TL (group) and DeepPHiC-ML (group) respectively. For DeepPHiC-ML (group), chromatin interactions in the group except tissue/cell type of interest are used to pretrain the shared feature extractor, chromatin interactions from tissue/cell type of interest are used to train the tissue-specific feature extractor. The integrated features from both the common features and private features will connect to the tissue-specific classifier for predicting tissue-specific chromatin interactions. For DeepPHiC-TL (group), chromatin interactions in the group except the tissue/cell type of interest is used to pretrain the common feature extractor, and the chromatin interactions from the tissue/cell type of interest is used to fine-tune the DeepPHiC-TL (group).

Consequently, for PE prediction, DeepPHiC-TL (group) outperforms DeepPHiC-TL in most cell types in H1 group (ME, MSC, NPC), and is comparable to DeepPHiC-TL in Brain group and Heart group except Left Ventricle (LV) (Figure 6A). In contrast, DeepPHiC-ML (group) has consistently comparable performance to DeepPHiC-ML (Figure 6C). For PP prediction, DeepPHiC-TL (group) is on par with DeepPHiC-TL in H1 group and Brain group but DeepPHiC-TL (group) is inferior to DeepPHiC-TL in Heart group (Figure 6B). One explanation is that the chromatin interactions in Heart group are few (e.g,CM (962), AO (788), LV (968), RV (723)). Therefore, the shared feature extractor may not be learnt accurately due to the small amount of labelled data even the cell types used are more relevant to the cell type of interest. Since Brain group has a much larger amount of chromatin interactions (e.g., HCmerge (2519), X5628FC (6224), the tissue/cell type-relevancy and the large sample size make the prediction performance comparable to using chromatin interactions from all tissues/cell types. Similarly, DeepPHiC-ML (group) is on par with DeepPHiC-ML in H1 group and Brain group but less favorable in Heart group (Figure 6D). Again, all methods follow a similar trend if the prediction performance are measured by AUPRC (Figure S4).

**Figure 6:**
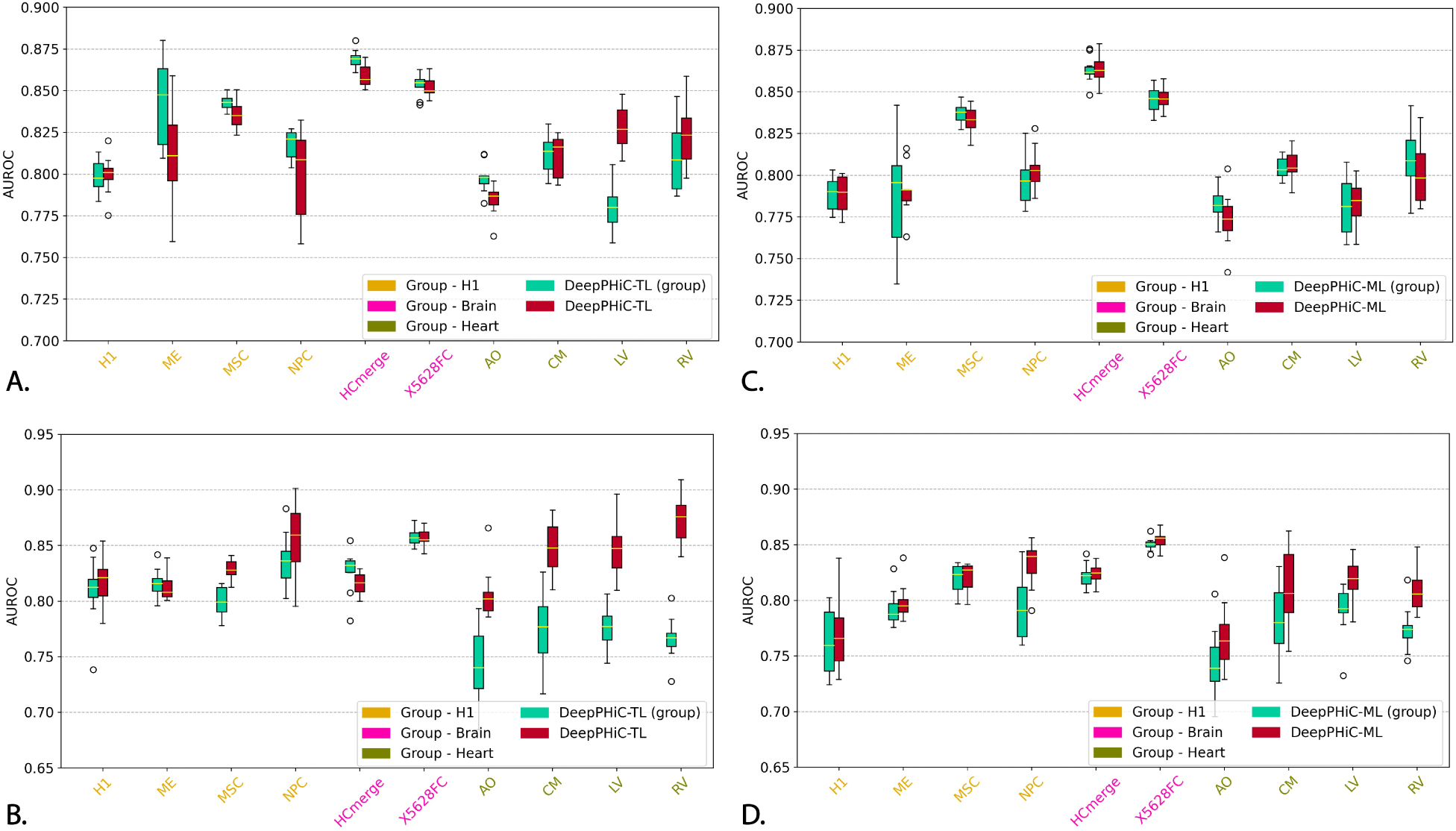
Compare DeepPHiC-TL and DeepPHiC-ML utilizing biologically relevant tissues/cell types and all tissues/cell types on predicting **A**. Promoter-enhancer interactions using DeepPHiC-TL and DeepPHiC-TL (group); **B**. Promoter-promoter interactions using DeepPHiC-TL and DeepPHiC-TL (group); **C**. Promoter-enhancer interactions using DeepPHiC-ML and DeepPHiC-ML (group); **D**. Promoter-promoter interactions using DeepPHiC-ML and DeepPHiC-ML (group). Each boxplot represents the AUCs in 10 experiments for each tissue/cell type. DeepPHiC-ML: DeepPHiC-ML uses all tissues/cell types. DeepPHiC-ML (group): DeepPHiC-ML uses biologically relevant tissues/cell types. DeepPHiC-TL: DeepPHiC-TL uses all relevant tissues/cell types. DeepPHiC-TL (group): DeepPHiC-TL uses biologically relevant tissues/cell types. Each boxplot represent the AUC in 10 experiments for each tissue.

**Figure 7:**
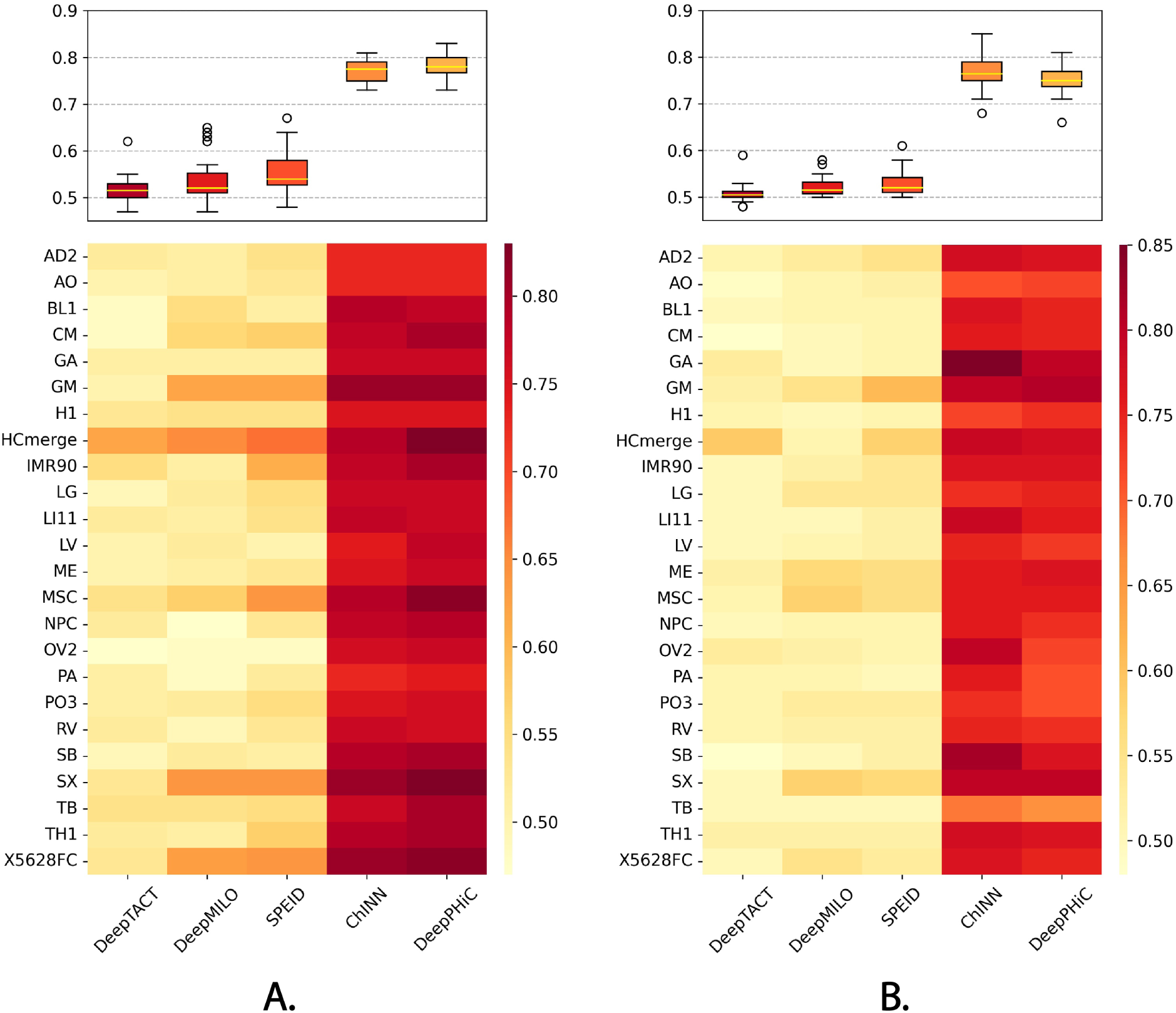
Comparing DeepPHiC and state-of-the-art deep learning models in “cross prediction”. **A**. Trained on PE and tested on PP (PE-to-PP) **B**. Trained on PP and tested on PE (PP-to-PE)

### 3.6 Comparing DeepPHiC and state-of-the-art deep learning models to cross predict promoter-enhancer interaction and promoter-promoter interaction

In the previous sections, we’ve demonstrated that DeepPHiC improves the prediction for both PE and PP. Here, we explore the possibility of “cross prediction”, where DeepPHiC is trained on PE and used to predict PP (PE-to-PP), and vice versa (PP-to-PE). Similarly, we include all aforementioned deep learning models in comparison and report the prediction performance in terms of AUC. As a result, for PE-to-PP, DeepPHiC and ChINN significantly outperform other approaches (DeepPHiC: 0.780; ChINN: 0.775; DeepTACT: 0.515; DeepMILO: 0.520; SPEID: 0.540). Similarly, for PP-to-PE, DeepPHiC and ChINN are also superior to other methods (DeepPHiC: 0.750; ChINN: 0.765; DeepTACT: 0.505; DeepMILO: 0.515; SPEID: 0.520). Therefore, “cross prediction” follows a similar trend as PE-to-PE and PP-to-PP. Moreover, the superiority of DeepPHiC and ChINN indicates that anchor distance is an important feature for “cross prediction”.

## 4 Discussion

In this work, we develop a multi-modal deep learning framework named DeepPHiC, which is based on the densely connected convolutional neural network, to predict promoter-centered 3D chromatin interactions such as promoter-enhancer interactions and promoter-promoter interactions by leveraging a comprehensive sets of Promoter Capture Hi-C datasets across multiple tissues/cell types. Different from existing approaches, DeepPHiC utilizes a comprehensive set of informative features, ranging from genomic sequence and epigenetic signal in anchors as well as anchor distance. We further demonstrate that using all three feature types together achieves better prediction performance than using arbitrary combinations of feature sets as model input. Moreover, we find that anchor distance is an informative feature, which indicates that anchors in promoter-centered interactions are closer to than random anchors in the genome.

We further develop two paradigms of DeepPHiC, which is based on multi-task learning (DeepPHiC-ML) and transfer learning (DeepPHiC-TL) respectively. The key idea of DeepPHiC-ML and DeepPHiC-TL lies on using chromatin interactions from other tissues to boost the learning for common feature space and thus fewer labelled data from targeted tissue is needed to train the tissue-specific prediction model, which in turn will improve the prediction for chromatin interactions in the targeted tissue. To demonstrate the effectiveness of the two paradigms of DeepPHiC, we compare both DeepPHiC-ML and DeepPHiC-TL to the baseline of DeepPHiC (DeepPHiC-Base), which is trained only using chromatin interactions in the targeted tissue, for both PE and PP prediction. Consequently, DeepPHiC-ML and DeepPHiC-TL outperforms DeepPHiC-Base in both PE and PP prediction, and DeepPHiC-TL achieves best prediction performance for both PE and PP prediction. Moreover, the advantage of DeepPHiC-ML and DeepPHiC-TL is more evident for tissues with scarce labelled data. As PP is much fewer than PE, DeepPHiC-ML and DeepPHiC-TL hold a more clear advantage in PP prediction than PE prediction. Due to the advantage of DeepPHiC-TL, we adopt DeepPHiC-TL as the default implementation when using DeepPHiC.

Furthermore, we compare DeepPHiC to state-of-the-art deep learning approaches with the similar purpose of predicting 3D chromatin interactions. As a result, DeepPHiC outperforms other competing approaches in predicting both PE and PP, which include SPEID and DeepMILO utilizing only genomic sequence in the anchors, ChINN using both anchor distance and genomic sequence in the anchors, and DeepTACT adopting genomic sequence and chromatin accessibility in the anchors. We find that DeepPHiC has the best performance and holds a clear advantage to other methods. Again, the improvement of DeepPHiC is more evident for tissues with a small amount of labelled data, especially for PP prediction. Moreover, we design a “cross prediction” strategy, where DeepPHiC is trained on PE and then used to predict PP, and vice versa. Still, we find a consistent trend that DeepPHiC and ChINN outperform other competing approaches.

Two important factors may affect the performance of DeepPHiC. One is the type of input features and the other is the tissues/cell types used for multi-task/transfer learning. Similar to ChINN, DeepPHiC incorporates the informative feature-“anchor distance” to improve the prediction performance. First, we perform an experiment to evaluate the robustness of DeepPHiC by removing anchor distance from the model input. Consequently, DeepPHiC without anchor distance still outperforms competing methods. Second, we evaluate the modified DeepPHiC-TL and DeepPHiC-ML by utilizing only biologically relevant tissues/cell types. As a result, we find that utilizing biologically relevant tissues/cell types will achieve comparable performance to using all relevant tissues/cell types when the number of chromatin interactions in biologically relevant tissues/cell types is reasonably large. However, the prediction performance of DeepPHiC-TL and DeepPHiC-ML declines when the number of chromatin interactions from biologically relevant tissues/cell types is limited.

## Acknowledgement

This work was supported by Indiana University Precision Health Initiative, Showalter Research Trust and National Institute of General Medical Sciences of the National Institutes of Health under Award Number R35GM142701 to LC.

## Notes

### Competing Interest Statement

The authors have declared no competing interest.

