## Supplemental File for "DeepPHiC: Predicting promoter-centered chromatin interactions using a novel deep learning approach"

### Supplementary materials for DeepPHiC: Predicting promoter-centered chromatin interactions using a novel deep learning approach

#### 1 Supplementary tables

| Tissue/Cell Type | Tissue/Cell Type Class | Abbr. | #PE (FDR<0.1) | #PP (FDR<0.1) |
|---|---|---|---|---|
| Adrenal gland | Primary Tissue | AD2 | 2556 | 5421 |
| Aorta | Primary Tissue | AO | 5224 | 788 |
| Bladder | Primary Tissue | BL1 | 4918 | 555 |
| Cardiomyocytes | Primary Tissue | CM | 7332 | 962 |
| Esophagus | Primary Tissue | EG2 | 20049 | - |
| Fat | Primary Tissue | FT2 | 56057 | - |
| Gastric tissue | Primary Tissue | GA | 1610 | 900 |
| GM12878+GM19240 Lymphoblastoid Cell Line | Primary Cell Line | GM | 4903 | 4446 |
| H1 Embryonic Stem Cell | Embryonic Stem Cell | H1 | 8178 | 917 |
| Hippocampus | Primary Tissue | HCmerge | 17434 | 2519 |
| Fibroblast cells | Primary Cell Line | IMR90 | 13217 | 4156 |
| Lung | Primary Tissue | LG | 1188 | 661 |
| Liver | Primary Tissue | LI11 | 5325 | 706 |
| Left Ventricle | Primary Tissue | LV | 1565 | 968 |
| H1-derived Mesendoderm Cell | Early Embryonic Lineages | ME | 1045 | 3142 |
| H1-derived Mesenchymal Stem Cell | Early Embryonic Lineages | MSC | 16379 | 4168 |
| H1-derived Neuronal Progenitor Cell | Early Embryonic Lineages | NPC | 3010 | 546 |
| Ovary | Primary Tissue | OV2 | 1105 | 187 |
| Pancreas | Primary Tissue | PA | 2594 | 396 |
| Psoas | Primary Tissue | PO3 | 1964 | 314 |
| Right Ventricle | Primary Tissue | RV | 1405 | 723 |
| Small Bowel | Primary Tissue | SB | 972 | 704 |
| Sigmoid colon | Primary Tissue | SG1 | - | 10338 |
| Spleen | Primary Tissue | SX | 22392 | 5866 |
| Trophoblast | Early Embryonic Lineages | TB | 83604 | 444 |
| Thymus | Primary Tissue | TH1 | 8993 | 1066 |
| Dorsorateral prefrontal cortex | Primary Tissue | X5628FC | 26084 | 6224 |

Table S1: Summary of promoter-enhancer and promoter-promoter interactions in 26 tissues/cell types

| Tissue/Cell Type | #PE | | #PP | |
|---|---|---|---|---|
| | Range | Median | Range | Median |
| Primary Tissue | 972 - 56057 | 3756 | 187 - 10338 | 788 |
| Primary Cell Line | 4903 - 13217 | 7332 | 962 - 4446 | 4156 |
| Embryonic Stem Cell | 8178 | 8178 | 917 | 917 |
| Early Embryonic Lineages | 1045 - 83604 | 9695 | 444 - 4168 | 1844 |

Table S2: Summary of promoter-enhancer and promoter-promoter interactions in four categories of tissues/cell types

| Tissue/Region | Roadmap Epigenomics (Chromatin Accessibility/Histone Modification |
|---|---|
| Adrenal gland | Pancreas Chromatin Accessibility |
| Aorta | Aorta H3K4me1 |
| Bladder | Fetal Kidney Chromatin Accessibility |
| Cardiomyocytes | Aorta H3K4me1 |
| Esophagus | Esophagus H3K4me1 |
| Fat | Adipose Tissue H3K27ac |
| Gastric tissue | Gastric Chromatin Accessibility |
| GM12878+GM19240 Lymphoblastoid Cell Line | Fetal Spleen Chromatin Accessibility |
| H1 Embryonic Stem Cell | H1 H3K4me3 |
| Hippocampus | Brain Hippocampus Middle H3K4me1 |
| Fibroblast cells | Breast Fibroblast Primary Cells H3K4me1 |
| Lung | Fetal Lung Right Chromatin Accessibility |
| Liver | Adult Liver H3K4me3 |
| Left Ventricle | Left Ventricle H3K4me1 |
| H1-derived Mesendoderm Cell | H1 Derived Mesenchymal Stem Cells H3K4me1 |
| H1-derived Mesenchymal Stem Cell | H1 Derived Mesenchymal Stem Cells H3K4me1 |
| H1-derived Neuronal Progenitor Cell | H1 Derived Neuronal Progenitor Cultured Cells H3K4me1 |
| Ovary | Fetal Ovary Chromatin Accessibility |
| Pancreas | Pancreas Chromatin Accessibility |
| Psoas | Psoas Muscle Chromatin Accessibility |
| Right Ventricle | Right Ventricle H3K4me1 |
| Small Bowel | Small Intestine Chromatin Accessibility |
| Sigmoid colon | Sigmoid Colon H3K4me1 |
| Spleen | Spleen H3K4me1 |
| Trophoblast | H1 BMP4 Derived Trophoblast Cultured Cells H3K4me1 |
| Thymus | Fetal Thymus Chromatin Accessibility |
| Dorsorateral prefrontal cortex | Fetal Brain Chromatin Accessibility |

Table S3: Summary of matched tissue/cell type-specific epigenetic data for pHi-C data

### 2 Supplementary figures

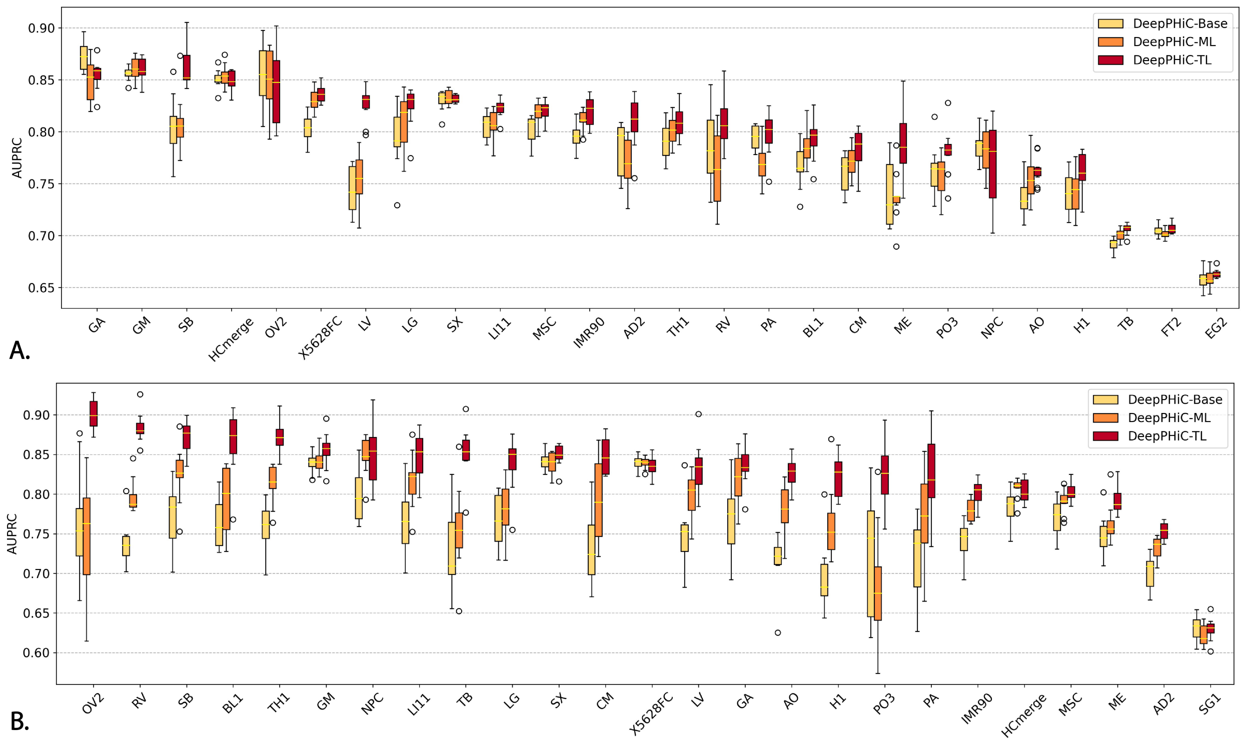


Figure S1: Comparison of three versions of DeepPHiC. **A.** promoter-enhancer interactions (PE) in 26 tissues/cell types and **B.** promoter-promoter interactions (PP) in 25 tissues/cell types. AUPRC are reported on 10 different experiments.

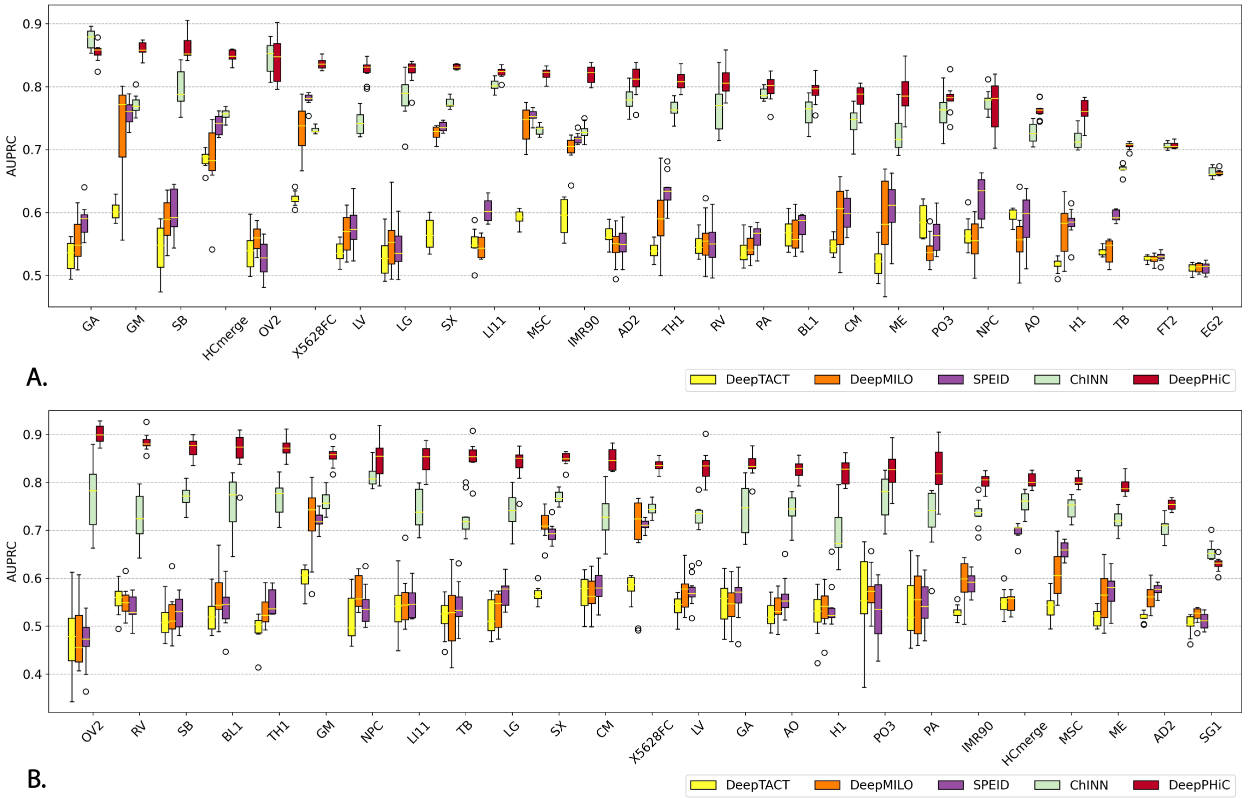


Figure S2: Comparison of DeepPHiC and state-of-the-art deep learning models on predicting **A.** Promoter-enhancer interactions (PE) and **B.** Promoter-promoter interactions (PP). Each boxplot represents the AUPRC in 10 experiments for each tissue.

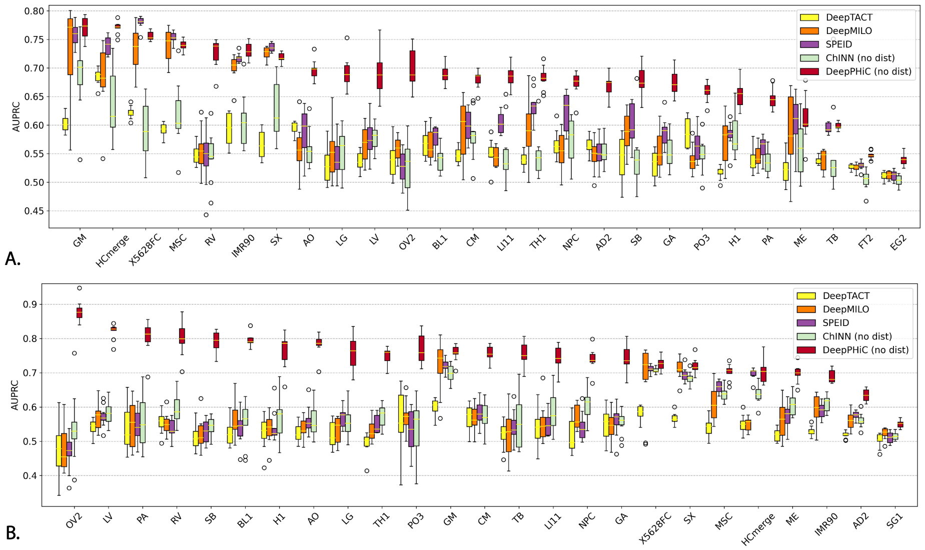


Figure S3: Compare DeepPHiC without distance (DeepPHiC (no dist))and ChINN without distance (ChINN (no dist)) to other deep learning models for predicting **A.** Promoter-enhancer interactions (PE) and **B.** Promoter-promoter interactions (PP). Each boxplot represents the AUC in 10 experiments for each tissue.

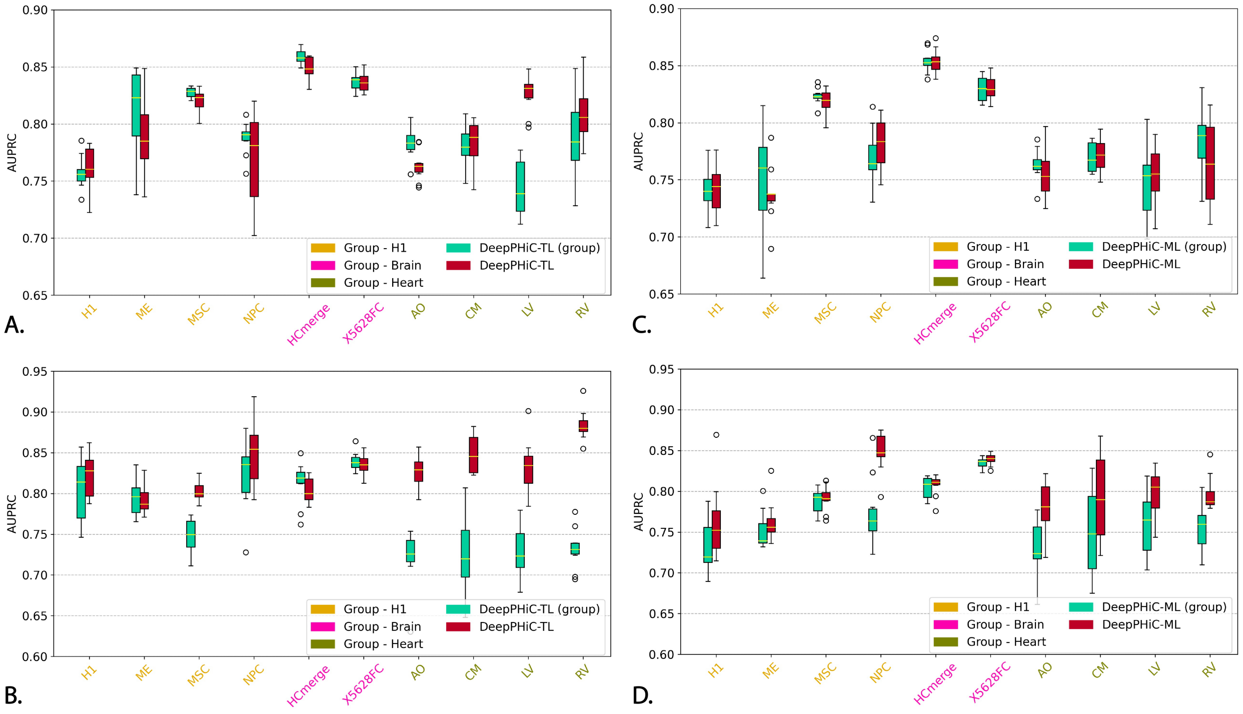


Figure S4: Compare DeepPHiC-TL and DeepPHiC-ML utilizing biologically relevant tissues/cell types and all tissues/cell types on predicting **A.** Promoter-enhancer interactions using DeepPHiC-TL and DeepPHiC-TL (group); **B.** Promoter-promoter interactions using DeepPHiC-TL and DeepPHiC-TL (group); **C.** Promoter-enhancer interactions using DeepPHiC-ML and DeepPHiC-ML (group); **D.** Promoter-promoter interactions using DeepPHiC-ML and DeepPHiC-ML (group). Each boxplot represents the AUC in 10 experiments for each tissue/cell type. DeepPHiC-ML: DeepPHiC-ML uses all tissues/cell types. DeepPHiC-ML (group): DeepPHiC-ML uses biologically relevant tissues/cell types. DeepPHiC-TL: DeepPHiC-TL uses all relevant tissues/cell types. DeepPHiC-TL (group): DeepPHiC-TL uses biologically relevant tissues/cell types.
